# Kallikrein-related peptidase 14 activates zymogens of membrane type matrix metalloproteinases (MT-MMPs) - a CleavEx library-based analysis

**DOI:** 10.1101/2020.04.23.057109

**Authors:** Katherine Falkowski, Ewa Bielecka, Ida B. Thøgersen, Oliwia Bocheńska, Karolina Płaza, Magdalena Kalińska, Laura Sąsiadek, Małgorzata Magoch, Aleksandra Pęcak, Magdalena Wiśniewska, Natalia Gruba, Magdalena Wysocka, Anna Wojtysiak, Magdalena Brzezińska-Bodal, Kamila Sychowska, Anastasija Pejkovska, Maren Rehders, Georgina Butler, Christopher M Overall, Klaudia Brix, Grzegorz Dubin, Adam Lesner, Andrzej Kozik, Jan J. Enghild, Jan Potempa, Tomasz Kantyka

## Abstract

Kallikrein-related peptidases (KLKs) and matrix metalloproteinases (MMPs) are secretory proteinases known to proteolytically process components of the extracellular matrix (ECM), thus modulating the pericellular environment in physiology and excessively in pathologies like cancer. However, the interconnection between these groups of proteases remains elusive. To test this hypothesis, we have developed a peptide library-based exposition system (**Cleav**age of **ex**posed amino acid sequences, CleavEx) aiming at investigating the potential of KLK14 to recognize and hydrolyze proMMP sequences specifically. Initial assessment of the library identified a total of ten MMP activation domain sequences which were validated by Edman degradation. The CleavEx analysis revealed that membrane-type (MT) MMPs are likely targeted by KLK14 for activation. Correspondingly, commercially available proMT-MMPs, namely proMMP14-17 were investigated *in vitro* and found to be effectively processed by KLK14. Again, the expected neo-N-termini of the activated MT MMPs were yielded and confirmed by Edman degradation. In addition, the productivity of proMMP activation was analyzed by gelatin zymography, which indicated the release of fully active, mature MT-MMPs upon KLK14 treatment. Lastly, MMP14 was shown to be processed on the cell surface by KLK14 using murine fibroblasts stably overexpressing human MMP14.

Herein, we propose KLK14-mediated selective activation of cell-membrane located MT-MMPs as an additional layer of their regulation within the ECM. As both, KLKs and MT-MMPs are implicated in cancer, the activation described herein may constitute an important factor in tumor progression and metastasis.

## INTRODUCTION

The extracellular matrix (ECM) consists of a complex array of locally secreted macromolecules interacting as a meshwork of proteins, glycoproteins, glycosaminoglycans, and proteoglycans [1]. Proteases present within the ECM are essential for turnover of structural proteins or core protein components. In addition, proteolytic enzymes are required for the activation of growth factors and other ligands of cell surface receptors. Thereby, proteolytic processing of ECM components and the molecules stored therein provides both, the structural organization and essential initial steps in cell-cell communication. By initiation of the signal transduction, proteases effectively regulate gene transcription eventually impacting on cell differentiation, proliferation, growth and apoptotic programs [2]. Proteolytic enzymes essential for ECM homeostasis predominantly belong to the families of a disintegrin and metalloproteinase (ADAMs) [3], matrix metalloproteinases (MMPs) [1], and tissue kallikrein-related peptidases (KLKs) [4]. The MMP family is considered especially important for ECM remodeling [5], [6]. In particular, the MMP proteases of the membrane-type subfamily (MT-MMPs) are essential in regulating cell migration during physiological wound healing [7], but are also implicated in cancer progression as they often involve in the initial steps of cancer cell metastasis [8]–[10].

It is corroborated that the activation of MT-MMPs happens intracellularly by means of furin-mediated processing such that the active forms of MT-MMPs decorate the cell surface right upon secretory release of cellular products by exocytosis. This pathway is not exclusive, as extracellular processing of proMT1-MMP (proMMP14) has been reported [11], [12]. About one third of all proMMPs activation domains contain an Arg/Lys-rich motif, compatible with furin specificity, but also targeted by serine proteases, particularly plasmin [13], [14] and transmembrane serine proteinases hepsin and TMPRSS2 [15].

The family of human kallikrein-related peptidases consists of a total of 15 different serine proteases with trypsin-like, chymotrypsin-like or mixed substrate specificity. Physiological roles of KLKs include regulation of cell growth and tissue remodeling [16]. Typically, KLKs and MMPs are coexpressed in many cell types and tissues [17]–[20], therefore understanding their interactions may provide a novel perspective for deciphering regulation of MMP activities in health and disease. A recent report has identified a pericellular network of KLK4, KLK14, MMP3, and MMP9 that are involved in initiation and promotion of prostate cancer cell metastasis [15], however the function of KLK14 in this complex was not identified.

Herein, we have employed a new unbiased technique CleavEx (**Cleav**age of **ex**posed amino acid sequences), which allowed us to test for the activation of MMP zymogen forms by KLK14. Our approach was based on the recombinant attachment of an MMP activation site sequence to the proteolytically stable HmuY protein. A full library of hybrid proteins displaying all 23 activation sequences of human proMMPs was expressed and purified. This library was then incubated with KLK14 and the generated products were subsequently analyzed to identify the potential processing. Library analysis was followed by investigation of KLK14-mediated processing of native proMMP14-17. Lastly, we identified the cell surface processing of proMMP14 by KLK14. To our knowledge, this is the first study examining the potential of KLK family member to activate the proMMP family.

## RESULTS

### CleavEx_proMMP_ library screening

Hybrid proteins representative of each individual MMP propeptide sequence were created such that they are covalently engineered onto the proteolytic-resistant bacterial protein HmuY [21]. The hybrid proteins additionally featured an attached N-terminal extension constituting a HisTag followed by the to-be-investigated sequence. This resulted in generating a library of 23 individual proteins, each exposing a sequence corresponding to the activation domain of the respective human prometalloproteinase. To verify the effectiveness of KLK14 in recognizing these peptide sequences, 25 ng of each hybrid protein was incubated with increasing concentrations of KLK14. The results were visualized by a HisTag-specific Western blot, where the loss of signal was indicative of the release of the HisTag *via* proteolytic removal of the fragment (Figure 1A). The stability of the fulllength HmuY protein was verified independently and no KLK14-mediated degradation was observed (data not shown). Among all 23 hybrid proteins tested in this manner, the sequences corresponding to proMMP11, proMMP14-17, proMMP21, proMMP24-25 and proMMP28 were recognized with highest effectiveness. This means to say that KLK14 at a concentration of 50 nM was able to completely remove the HisTag from the respective hybrid proteins with HmuY. Note that CleavEx_proMMP23_ was only partially cleaved at 50 nM KLK14, but completely removed when 250 nM KLK14 was used.

**Figure 1:**
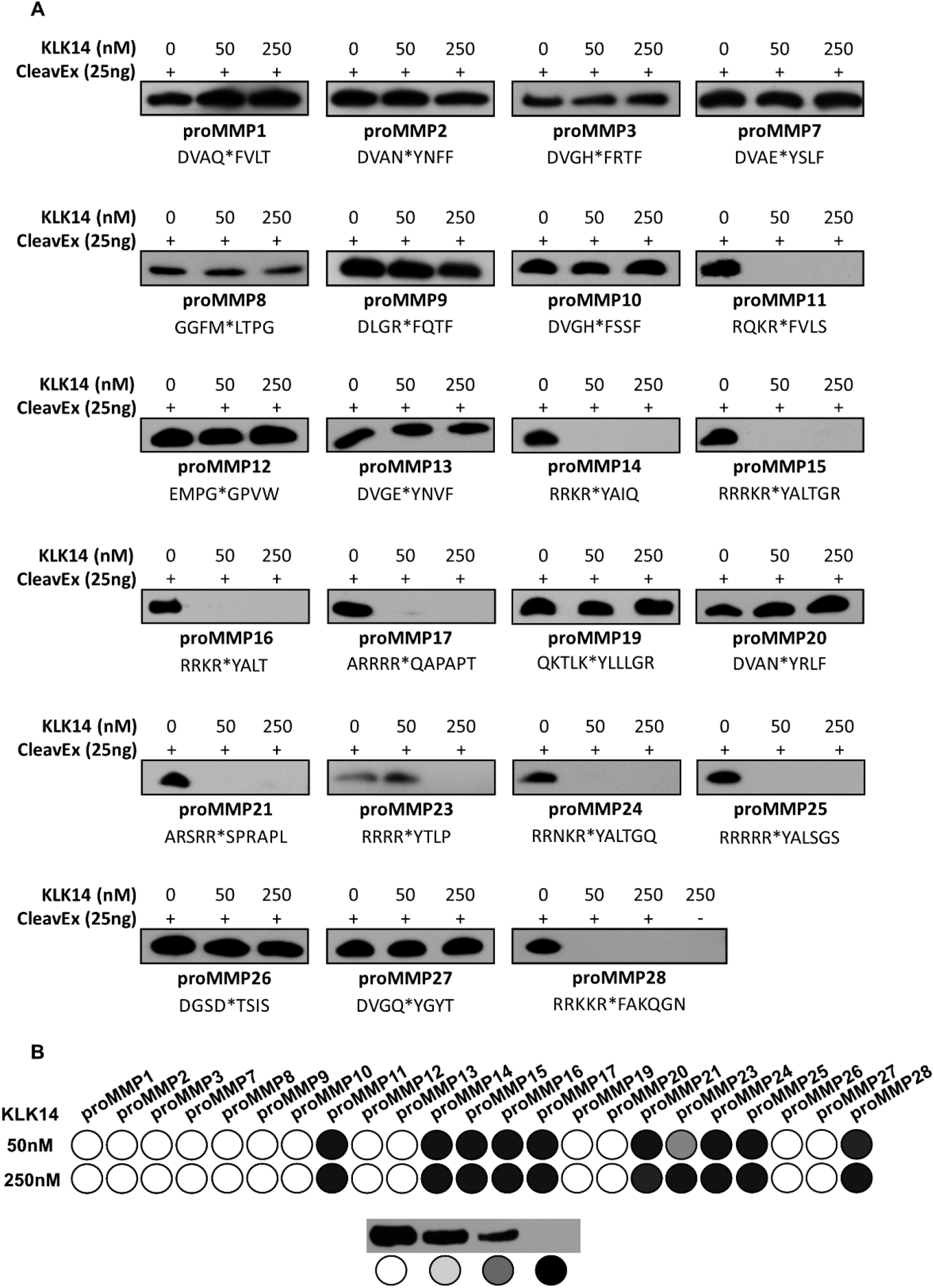
Effect of KLK14 on the CleavEx_proMMP_ library. **(A)** Western blot analysis of each 25 ng CleavEx_proMMP_ protein incubated with 50 and 250 nM KLK14 after 1 hour at 37°C. Each fusion protein with its respective activation sequence is listed with the native site of hydrolysis indicated by an asterisk. **(B)** Schematic representation of the CleavEx_proMMP_ library with shading based on the scoring of Western blots in panel A; white – no hydrolysis, light grey – partial hydrolysis, grey – moderate hydrolysis and black – full hydrolysis.

To visualize the efficiency of cleavage within the sequence motives of different CleavEx_proMMPs_ by KLK14, the Western blot results were scored and presented as a grey scale map (Figure 1B). The data indicated a striking selectivity of KLK14 towards particular proMMP sequences. To determine which peptide bonds were hydrolyzed within the activation domains, and to exclude the possibility of HmuY cleavage by KLK14, N-terminal sequencing of each hydrolyzed product of the CleavEx_proMMP_ reactions was performed. The determined N-termini nearly uniformly confirmed the expected P1 recognition site for KLK14 (Table 1). All of the CleavEx hybrid proteins except for CleavEx_proMMP21_, were hydrolyzed at the activation-specific location expected for KLK14.

**Table 1:**
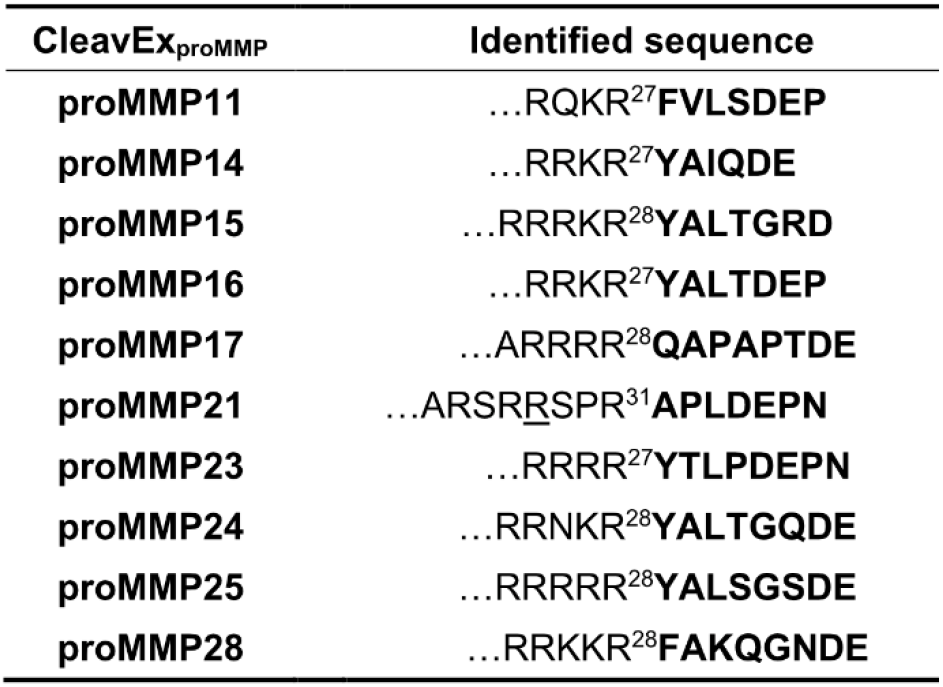
Identification of the KLK14 hydrolysis sites within the CleavEx_proMMP_ protein. CleavEx_proMMP_ fusion proteins were separated using SDS-PAGE and electrotransferred for N-terminal sequencing. Identified sequences are represented in the bold font and the underscore denotes where the location of the expected activation cleavage P1-P1’ in the proMMP-derived sequence.

### Processing of recombinant proMMPs by KLK14

Scanning of the hybrid protein library allowed for the selection of proMMPs that are likely activated by KLK14 *in situ.* To verify this contention, commercially available proMMP14, proMMP15, proMMP16 and proMMP17 were analyzed for KLK14-mediated activation *in vitro.* Firstly, we investigated the concentration requirements of KLK14-dependent hydrolysis of each proMMP. To this end, each proform was incubated with increasing concentrations of KLK14 for 1 hour (approx. from 1:65 to 1:5 KLK14:MMP molar ratio) and was then subjected to SDS-PAGE analysis. During incubation, the pan-specific MMP inhibitor batimastat was present in the reaction mixtures to prevent autoactivation of the proMMPs. ProMMP2 which was unaffected by KLK14 in the CleavEx system served as the negative control (Figure 2A). In parallel, furin was also analyzed with both, proMMP2 and proMMP14 as a negative and positive control, respectively (Figure 2B and D). All four proMMPs tested were effectively cleaved in the presence of KLK14 even at the highest substrate:enzyme molar ratios with discrete bands at molecular masses corresponding to the loss of the ~10 kDa propeptides. Moreover, the extent of proteolysis progressed in a concentrationdependent manner with 50 nM KLK14 (approx. 1:32 KLK14:MMP14 molar ratio) being sufficient to completely process proMMP14 (Figure 2C). Interestingly, in contrast to furin-mediated activation of MMP14, KLK14 incubation did not result in the accumulation of the ~12 kDa prodomain (Figure 2C and D). Higher concentrations of KLK14 were required for the full processing of other analyzed proMMPs. Nonetheless, 100 nM KLK14 was sufficient to process proMMP15-17 (approx. 1:16 KLK14:MMP15-17 molar ratio) and the hydrolyzed products remained stable up to 250 nM KLK14 (Figure 2E–G). Collectively, these results indicate that KLK14-mediated proMMP processing released stable, processed forms of MMP14-17.

**Figure 2:**
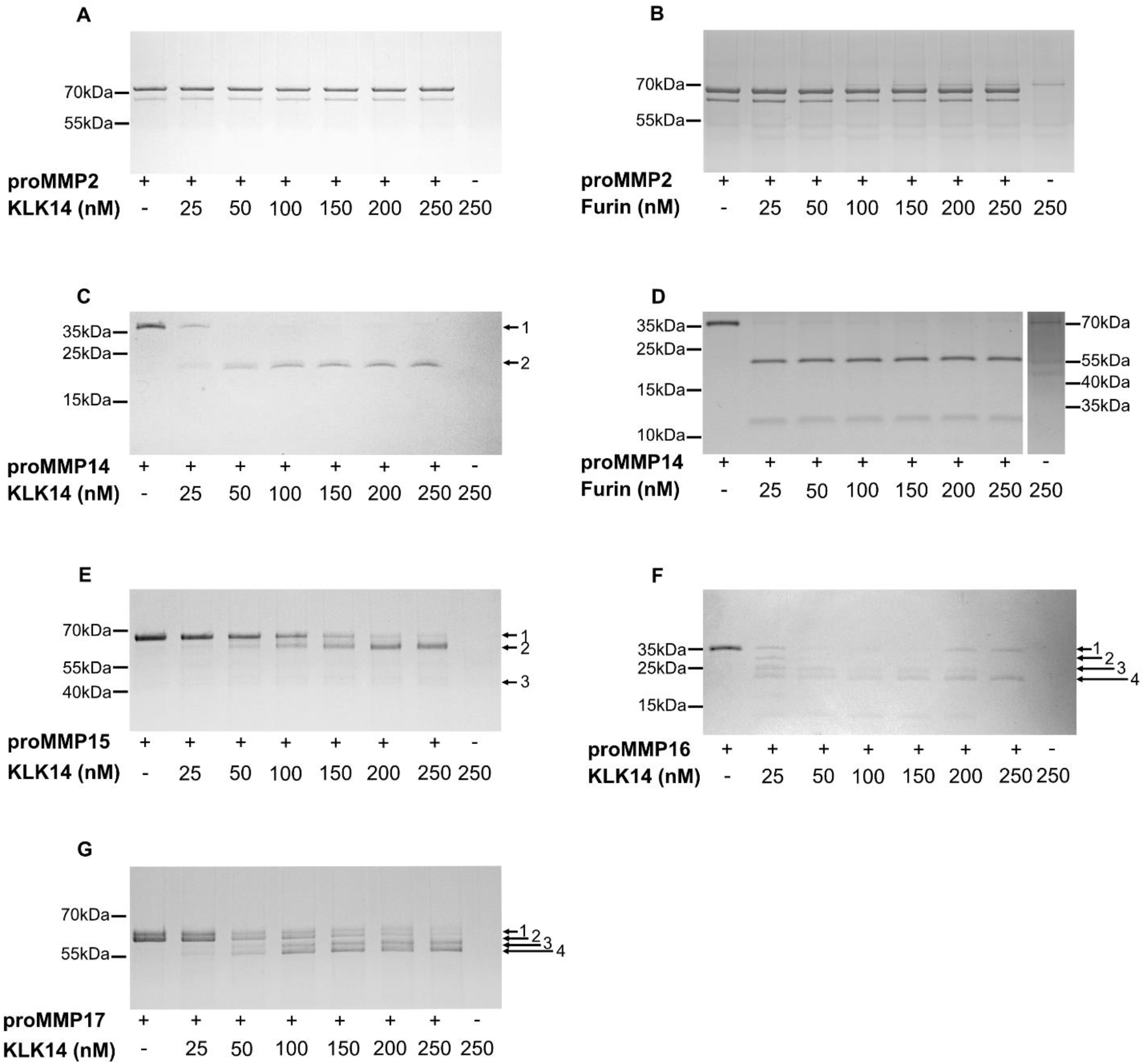
Concentration-dependent processing of proMMPs by KLK14. Each respective proMMP was incubated with increasing concentrations of KLK14 **(A, C, E-G)** or furin **(B, D)** for 1 hour at 37°C in the presence of 5 μM batimastat, the reaction products were analyzed by SDS-PAGE and visualized with Coomassie staining. Bands denoted with arrows were identified by N-terminal sequencing using Edman degradation (Table 2).

In addition to the products with the expected 10 kDa shift, several other transient forms were observed and subjected to Edman sequencing for unambiguous identification. The N-terminal analysis allowed for the assignment of the cleavage sites within the proform sequence (Table 2). First, the N-termini of all of the intact proforms, denoted as band 1, were confirmed as corresponding with the reported sequences of proMMPs, providing an internal quality control. The single product, band 2_MMP14_, generated by KLK14-mediated proMMP14 processing corresponded to the expected RRKR^111^→YAIQ sequence, previously identified as the activation site [11]. Similarly, KLK14-mediated proMMP15 hydrolysis resulted in the accumulation of band 2_MMP15_, *PG*KR^131^→YALT, consistent with the expected N-terminal sequence of the mature MMP15 [22]. Of note, despite the amino acids 128 and 129 being modified by the manufacturer from native arginine and arginine to proline and glycine, as indicated by italics above, KLK14 was able to generate the expected MMP15 active form. Furthermore, an additional product denoted as band 3_MMP15_ was detected to be DLRG^298^↓IQQL. Surprisingly, this product did not correspond to the expected KLK14 preference and rather was consistent with MMP15 specificity[23]. ProMMP16 processing lead to the formation of the KLK14-produced mature form, as band 4_MMP16_ corresponding to the anticipated RRKR^119^↓YALT hydrolysis. N-terminally truncated proforms were detected as band 2_MMP16_ and 3_MMP16_, however only the N-terminal of band 3_MMP16_ (KKPR^100^↓CG) corresponded to the expected KLK14 specificity while band 2_MMP16_ (ALAA^75^↓MQ) was consistent with MMP16 specificity [23]. ProMMP17 processing resulted in the generation of 4 bands. The first two bands, 1_MMP17_ and 2_MMP17_, observed in the untreated sample, were detected with identical N-termini A^39^PAPA, indicating the presence of differences in the C-terminal region of these proforms. KLK14 was able to process both proforms which resulted in the same sequential motif of TQAR^122^↓RRPQ (bands 3_MMP17_ and 4_MMP17_), three residues upstream of the expected native activation site.

**Table 2:**
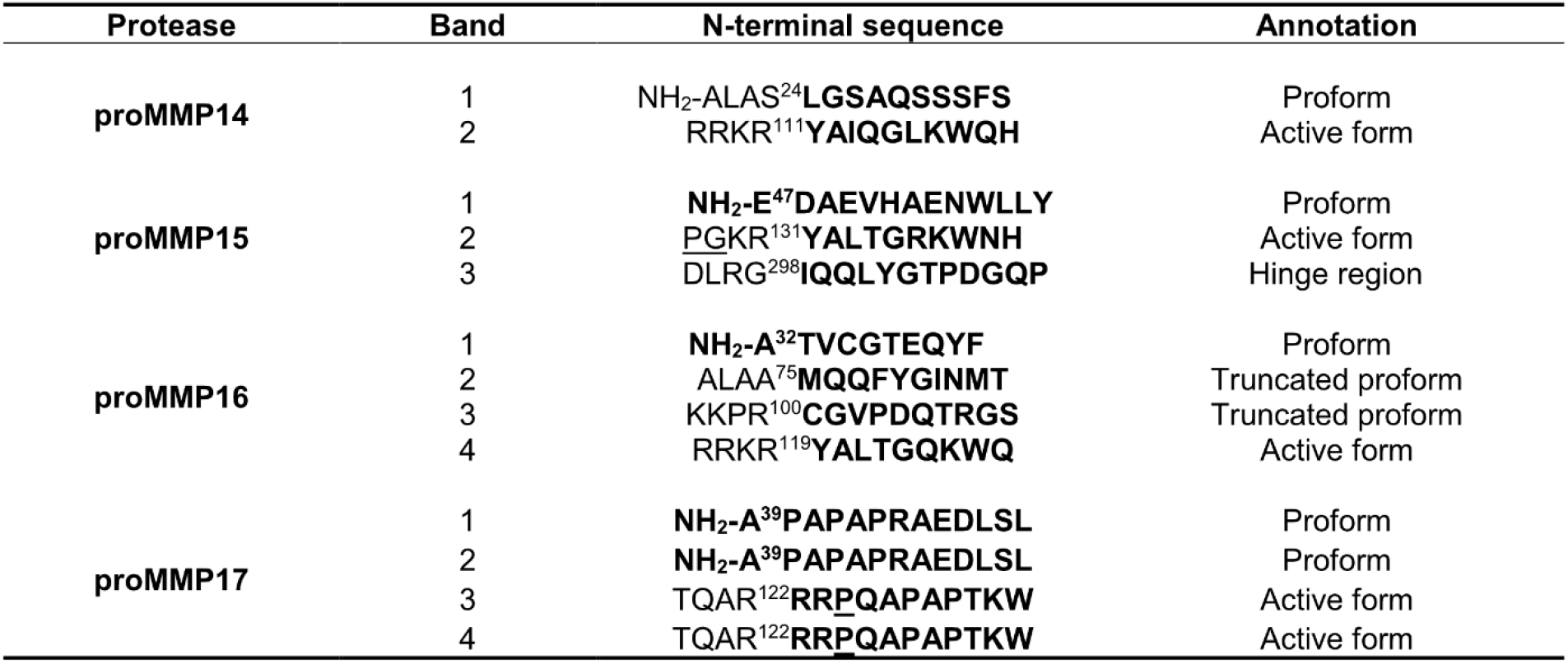
N - terminal identification of KLK14-mediated processing of recombinant proMMPs. The KLK14 hydrolysis product sequences were analyzed by N-terminal sequencing using Edman degradation. The bold font denotes the amino acid sequences identified. The underscored residues represent changes to the native protein sequence, as reported by the manufacturer (R&D Systems). KLK14 recognized the sequence 3-aa upstream of the native MMP17 activation site, likely because the native site was modified by the manufacturer. All residues are numbered according to the Uniprot reported sequence of the full-length proteins. Bands are labeled according to the notation explained at figure 2.

| Protease | Band | N-terminal sequence | Annotation |
| --- | --- | --- | --- |
| proMMP14 | 1 | NH<sub>2</sub>-ALAS<sup>24</sup>**LGSAQSSSFS** | Proform |
|  | 2 | RRKR<sup>111</sup>**YAIQGLKWQH** | Active form |
| proMMP15 | 1 | **NH<sub>2</sub>-E<sup>47</sup>DAEVHAENWLLY** | Proform |
|  | 2 | PGKR<sup>131</sup>**YALTGRKWNH** | Active form |
|  | 3 | DLRG<sup>298</sup>**IQQLYGTPDGQP** | Hinge region |
| proMMP16 | 1 | **NH<sub>2</sub>-A<sup>32</sup>TVCGTEQYF** | Proform |
|  | 2 | ALAA<sup>75</sup>**MQQFYGINMT** | Truncated proform |
|  | 3 | KKPR<sup>100</sup>**CGVPDQTRGS** | Truncated proform |
|  | 4 | RRKR<sup>119</sup>**YALTGQKWQ** | Active form |
| proMMP17 | 1 | **NH<sub>2</sub>-A<sup>39</sup>PAPAPRAEDLSL** | Proform |
|  | 2 | **NH<sub>2</sub>-A<sup>39</sup>PAPAPRAEDLSL** | Proform |
|  | 3 | TQAR<sup>122</sup>**RRPQAPAPTKW** | Active form |
|  | 4 | TQAR<sup>122</sup>**RRPQAPAPTKW** | Active form |

To follow up on the concentration analysis, a time-course experiment was employed to detect product accumulation by SDS-PAGE. Specific time points were analyzed for 3 hours with either 50 nM KLK14 (proMMP14) or 100 nM KLK14 (proMMP15, proMMP16 and proMMP17) to observe the processing of each respective proMMP (Figure 3). Within 5 minutes, the stable product, with the molecular weight corresponding to the active MMP14 form began to appear and near complete processing was observed after 3 hours, resulting in the accumulation of the mature MMP14 form (Figure 3A). In contrast, MMP15 degradation progressed more slowly with the band corresponding to the processed active form appearing vaguely after 30 min. Again, the product accumulated as a stable band, yet proMMP15 was not fully processed even after 3 hours (Figure 3B). On the other hand, proMMP16 products accumulated already within 5 minutes. All four bands identified by N-terminal sequencing appeared early, illustrating the efficient processing by KLK14. The fully matured active form appeared as the main band after 1 hour of reaction and remained stable for the whole 3 hours of incubation (Figure 3C). Again, similarly to proMMP15, proMMP17 maturation was not as efficient, as lower-molecular weight bands started to appear after 2 hours of reaction and complete processing of the proform could not be observed during the incubation (Figure 3D).

**Figure 3:**
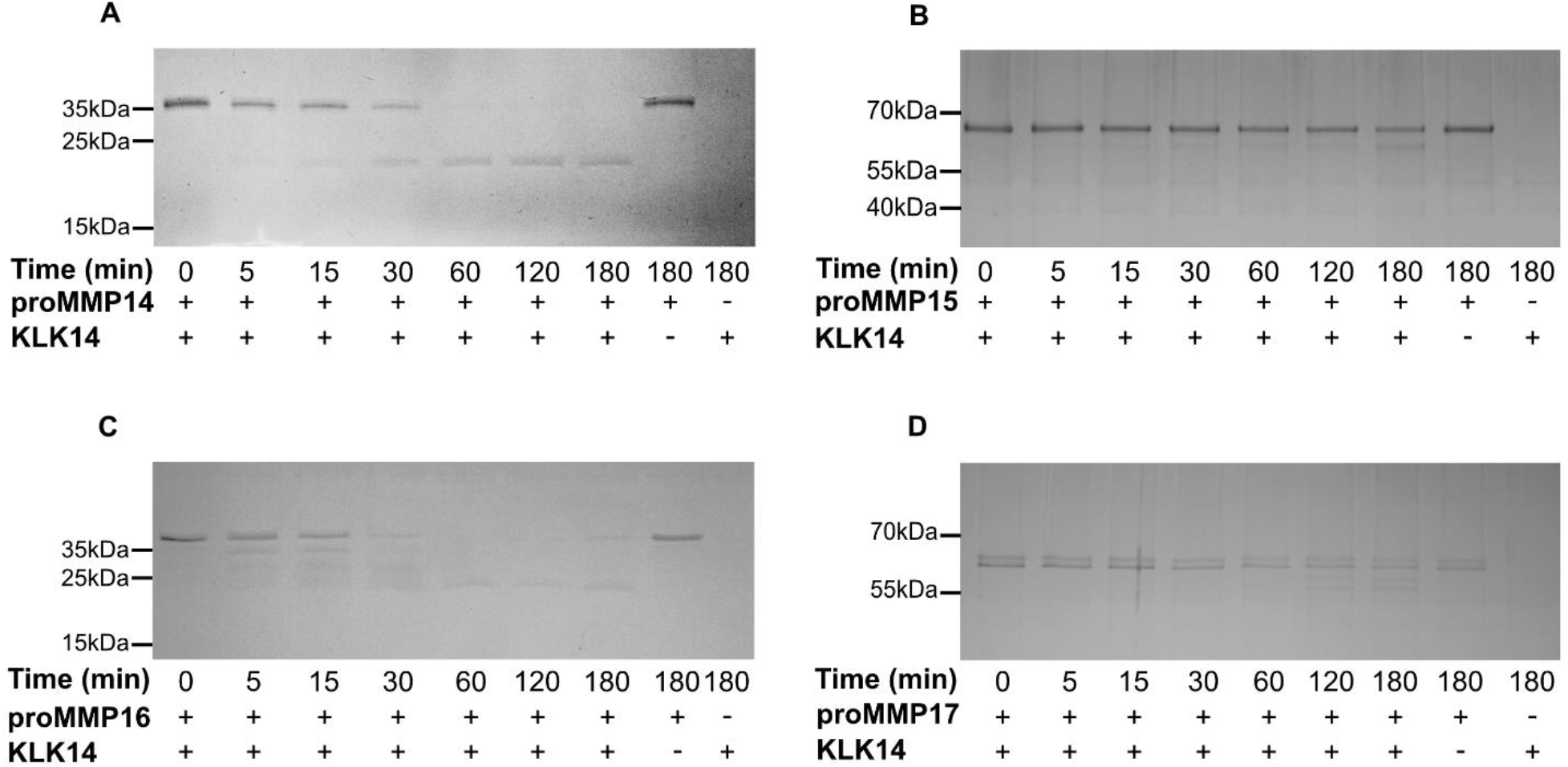
Recombinant human proMMP14,15, 16 and 17 are processed by KLK14 in a time dependent manner. **(A-D)** Respective proMMPs were incubated with KLK14 for a total of 3 hours at 37°C in the presence of 5 μM batimastat. The reaction products at indicated timepoints were analyzed by SDS-PAGE and visualized via Coomassie staining.

### Functional activation of proMMPs by KLK14

Each analyzed proMMP was activated with 50, 100 or 250 nM KLK14 for an hour at 37°C followed by a gelatin zymography. No KLK14 activity was detected under experimental conditions, as evident by the lack of clear bands in the KLK14 control lanes (Figure 4A-E). As previously mentioned, proMMP2 was used as a negative control. Regardless of proMMP2 alone displaying residual activity on the zymogram gel, KLK14 treatment resulted in no observable processing after 250 nM KLK14 incubation (Figure 4A). In contrast, gelatinolytic activity and the expected molecular mass shifts were observed for proMMP14 – 16. At both 50 and 100 nM KLK14, proMMP14 was processed to its active form with no difference in band intensity at either concentration (Figure 4B), indicating KLK14 concentrations equal or below 50 nM as sufficiently efficient convertase. Interestingly, a portion of active MMP15 was observed with no treatment of KLK14 which suggested self-activation during the 1-hour incubation (Figure 4C). Furthermore, KLK14 treatment of proMMP15 resulted in enhanced activation and active band accumulation at the expected molecular weight of around 55 kDa at 100 nM KLK14, which intensity slightly decreased at 250 nM KLK14. In addition, lower molecular mass bands (~35 kDa) were observed at both KLK14 concentrations, also displaying gelatinolytic activity. Part of these fragments likely corresponded to the DLRG^298^↓IQQL truncation identified from N-terminal sequencing as well as MMP15 self-degraded products since no MMP inhibitor is present in the activity assay. As expected, proMMP16 was activated and its gelatinolytic activity was stable at both KLK14 concentrations (Figure 4D). Lastly, there was no observable proMMP17 gelatinolytic activity despite the expected molecular mass shift of its mature form, visible in the zymogram gel (Figure 4E). MMP17 is unlikely acting as a gelatinase, which is consistent with some previous reports of others [24].

**Figure 4:**
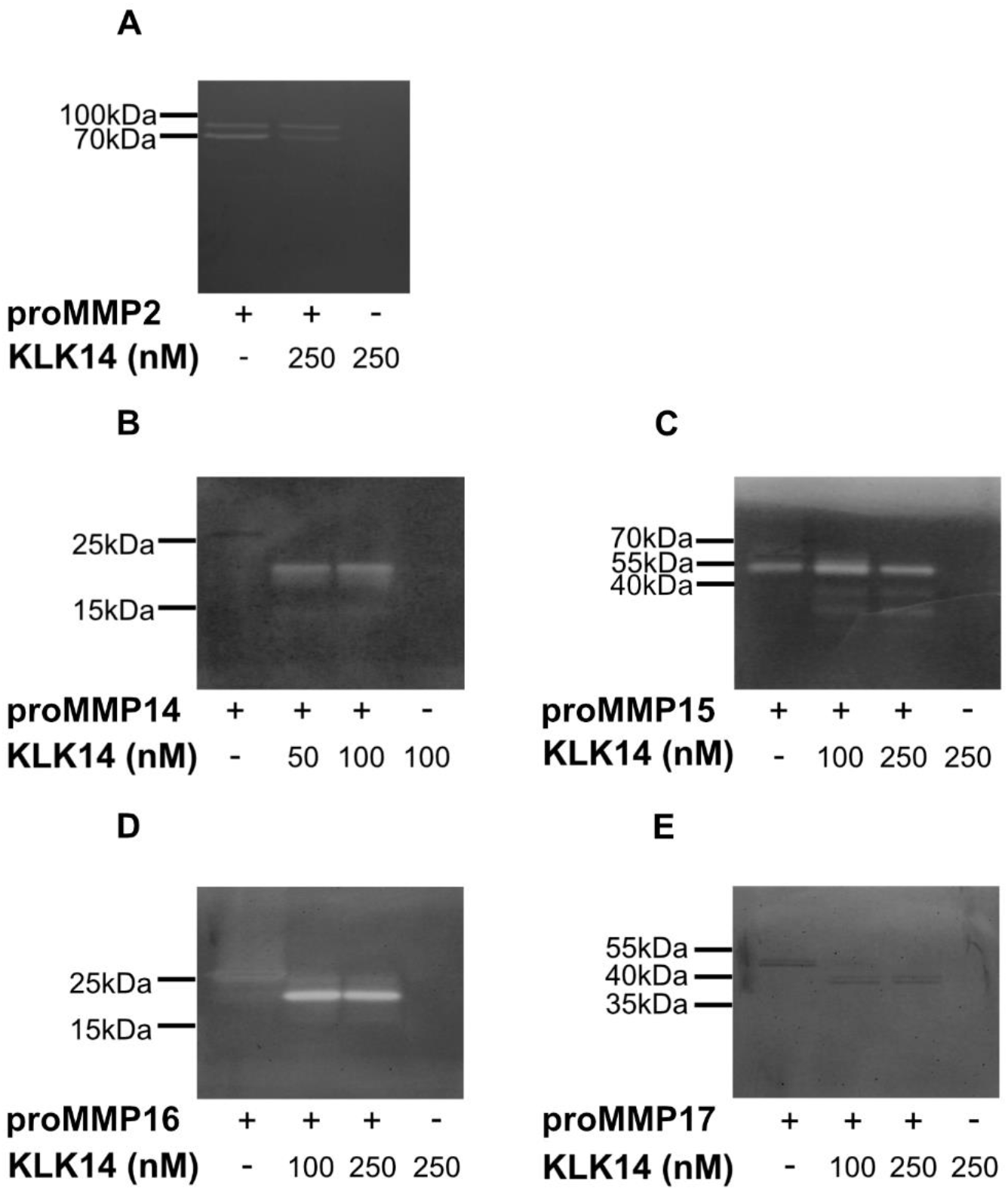
Gelatin zymography of proMMPs by KLK14-mediated processing. Activation of proMMPs by KLK14 results in a fully functional mature enzyme. Each proMMP was incubated with the indicated concentrations of KLK14 for an hour at 37°C. The reaction was stopped by the addition of KLK14-specific inhibitors and the reaction mixture was analyzed by SDS-PAGE, followed by a zymogram with gelatin as a substrate. The proMMP2 **(A)** negative control was not activated. ProMMP14 **(B)**, proMMP15 **(C)** and proMMP16 **(D)** were activated whereas proMMP17 **(E)** did not show hydrolysis of gelatin, yet a shift corresponding to the loss of the profragment was observed (note that an amino acid substitution was introduced in proMMP17 by the manufacturer (R&D Systems)).

### Comparison of functional activation of proMMP14 by KLK14 and furin

Native recombinant proteins illustrated that furin and KLK14 have similar proMMP14 activation efficiency. Therefore, to confirm the equivalence of KLK14 and furin processing, increase in proMMP14 activity on a synthetic substrate was observed in parallel upon treatment with these convertases. Observed activity of KLK14/furin-treated MMP14 increased evenly for 30 minutes. However, after an hour, the KLK14-mediated activation lead to 2-fold higher MMP14 activity, when compared to furin. This difference remained consistent for the remaining 180 minutes of hydrolysis. As expected, neither KLK14 or furin had no observable activity on the MMP14 fluorescent substrate (Figure 5).

**Figure 5:**
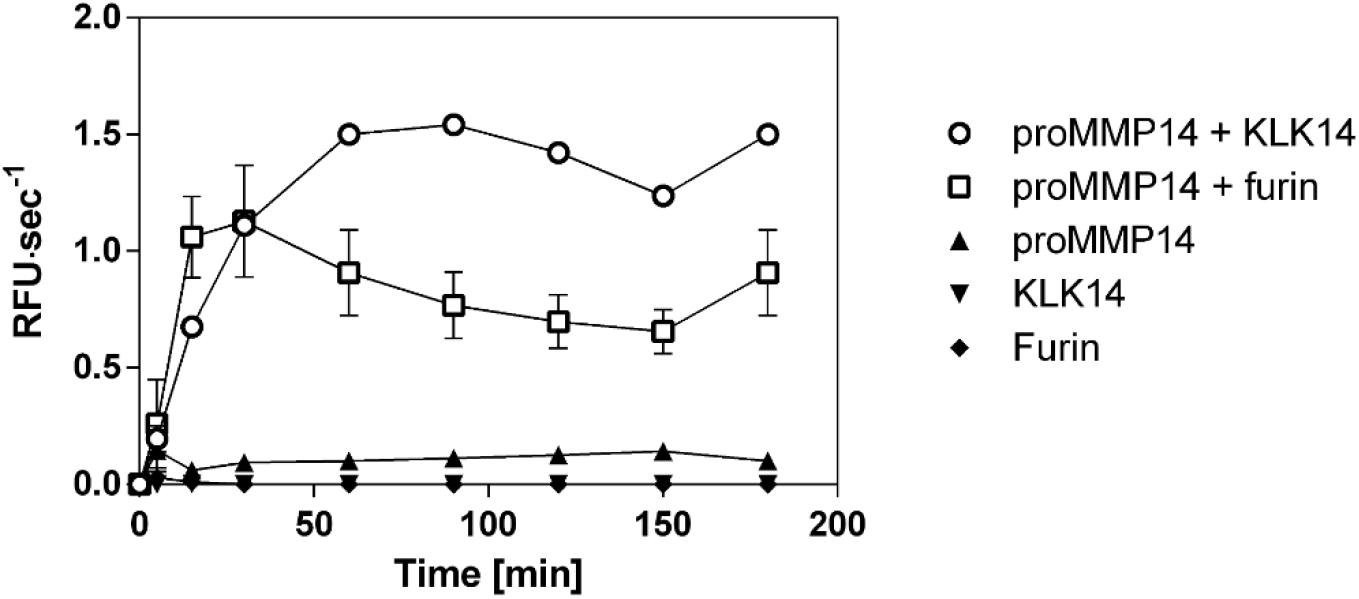
Comparison of recombinant human proMMP14 activation by KLK14 and furin as a functional peptidase. ProMMP14 (10 nM) was incubated with KLK14 (3 nM) and furin (3 nM) in the presence of a fluorogenic substrate Mca-KPLGL-Dpa-AR-NH_2_. The final concentration of the substrate was 10 μM in the MMP reaction buffer (50 mM Tris, 3 mM CaCl2, 1 μM ZnCl2, pH 8.5). Hydrolysis was recorded for 180 minutes at 37°C.

### Cell surface processing of proMMP14 by KLK14

A key step in demonstrating whether KLK14 activates proMMP14 at the cell surface is to validate whether it can occur in a cellular system. For this, murine embryonic fibroblasts stably expressed with human MMP14 (MT1-MMP) were grown to obtain a cell density of 1 million cells/well. Cells were treated with active KLK14 and furin along with appropriate controls and surface protein biotinylation was performed, followed by immunoprecipitation of the biotinylated proteins and a MMP14-specific Western blot. The specific bands for MMP14 were seen with a corresponding ~63 kDa and ~58 kDa molecular mass (Figure 6) and were interpreted as proMMP14 and active MMP14, respectively, in accordance with the previous records [25], [26]. Upon cell stimulation with 50 nM KLK14 for 30 min, an increase in intensity of the 58 kDa band was observed. Furthermore, a processed MMP14 form was detected at a slightly lower molecular mass of ~56 kDa. Interestingly, furin did not facilitate an increase in intensity in any of the bands, nor it produced any lower-molecular weight products, detectable on Western blot.

**Figure 6:**
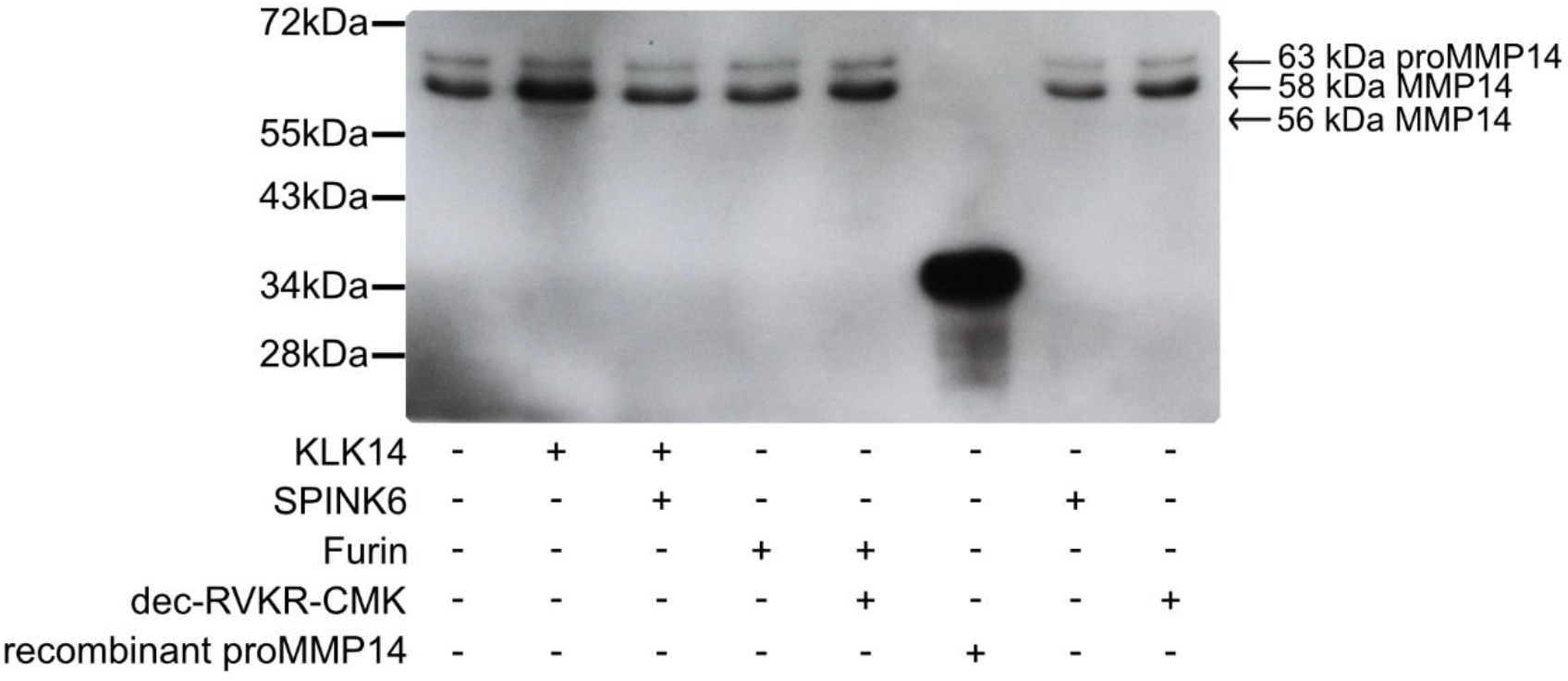
Processing of cell surface proMMP14 by KLK14. Murine embryonic fibroblasts stably expressing human MMP14 (MT1-MMP) were treated with KLK14 and furin as well as these enzymes inhibited then biotinylated followed by immunoblotting using an anti-MMP14 antibody. Each sample contained the 63 kDa proMMP14 form whereas an increase in the active 58 kDa MMP14 form was observed after KLK14 incubation. Additionally, a lower molecular weight MMP14 form at 56 kDa was detected only in the KLK14 treated sample. SPINK6 (KLK14 specific inhibitor) and dec-RVKR-CMK (furin inhibitor).

## DISCUSSION

Matrix metalloproteinases are essential enzymes in the maintenance of tissue homeostasis, namely by regulation of its self-reorganization, through growth factor activation and processing, as well as involvement in the regulation of immune cell functions [27]. Herein, we present biochemical evidence of selective activation of membrane-type MMPs by the kallikrein-related protease family member KLK14.

Our assessment is based on an unbiased approach, where all proMMP-specific activation sequences were displayed utilizing a hybrid protein system, allowing for easy production, purification and fragmentation analysis. The described herein CleavEx (**Cleav**age of **ex**posed amino acid sequences) system facilitated a rapid selection of KLK14-specific candidate proMMPs for further analysis with recombinant proteins. The carrier protein, HmuY, is produced by the bacterium responsible for periodontitis, *Porphyromonas gingivalis,* which uses an arsenal of proteolytic enzymes as virulence factors [28]. Stability studies indicate an unsurpassed resistance of the HmuY protein to proteolytic digestion by trypsin and bacterial proteinases alike [21] and suggest enhanced stability *in situ* (i.e. in inflamed tissue). Furthermore, structural studies revealed that the N- and C-termini of HmuY are exposed to the solvent thus, are easily accessible, while the overall globular structure of HmuY ensures good solubility and ease of folding when produced recombinantly [21]. The N-terminal HisTag serves as an affinity label for purification and additionally as a degradation-specific tag for follow-up of peptide bond hydrolysis, even in complex protein mixtures. Previously, a similar approach, but using a modified fibroblast growth factor-1 (FGF-1), was implemented by another group in the analysis of a proKLK cascade activation [29], [30] and MMP-dependent activation of KLKs [31]. Yet, we believe the choice of HmuY used herein to be superior. The approximate 25 kDa molecular masses of the CleavEx fusion proteins and the ~2.5 kDa N-terminal fragment removal allowed for an easy follow-up of the reaction products by SDS-PAGE and Western blot in comparison to the FGF-1-based system, characterized by a ~1kDa shift only.

Out of all 23 sequences, a total of 10 CleavEx_proMMP_ fusion proteins were hydrolyzed by KLK14, specifically the ones corresponding to membrane-type MMPs (MMP14-17 and MMP24-25), stromelysin-3 (MMP11), MMP21, femalysin (MMP23) and epilysin (MMP28) activation sequences. Importantly, CleavEx_proMMP_ hydrolysis by KLK14 was specific and limited to sequences derived from a subset of metalloprotease zymogens. Strikingly, the majority of all proMMPs recognized by KLK14 in the CleavEx analysis, cluster together constituting the group of membrane-type (MT) MMPs. This subgroup of the MMP proteinases is bound to the plasma membrane, either by type I transmembrane domains (MMP14-16, MMP24) or by means of phosphatidylinositol-anchors attached to the protein chains (MMP17 and MMP25) [18]. Therefore, recognition by KLK14 reflects not only the similarities in the activation domain of MT-MMPs, but also the common evolutionary ancestry and functional homology of the processed proteinases. Together, these results indicate that the group of cell-surface proMMPs can be nearly exclusively targeted by KLK14-mediated activation (Figure 7).

**Figure 7:**
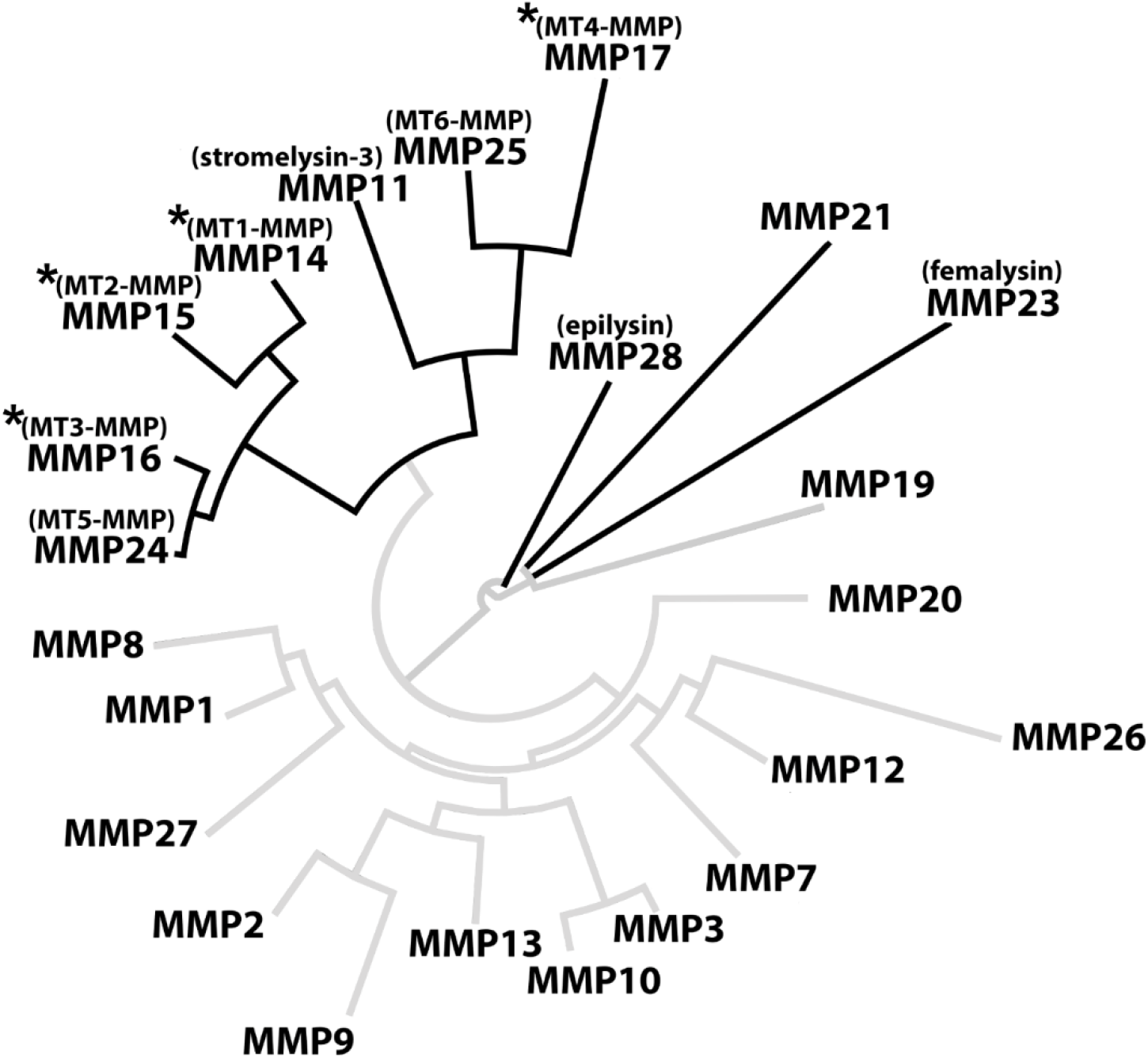
KLK14-activated MMPs cluster together in phylogenetic analysis. The full sequences of all human MMPs were obtained from the uniport database and analyzed in BioEdit using built-in multiple alignment ClustalW algorithm. The resulting alignment was visualized using an online tool (www.phylogeny.fr). The full black branches represent the MMPs in which the profragment activation sequence was hydrolyzed by KLK14. An asterisk denotes functionally activated MMPs as verified in vitro. A hashtag denotes that N-terminal sequencing of CleavEx_proMMP21_ was determined three amino acids downstream of the expected P1 activation site.

Commercially available recombinant proMMPs were investigated to confirm the initial CleavEx_proMMP_ analysis. KLK14 was an effective convertase for proMMP14 and proMMP16 but less-efficiently, for proMMP15 and proMMP17 in a time dependent manner. N-terminal sequencing of the concentration-dependent KLK14-released products identified the expected hydrolysis sites for proMMP14, proMMP15 and proMMP16, respectively, indicating native-like processing of these enzymes by KLK14 [11], [22]. ProMMP14 processing by KLK14 and furin revealed a single band corresponding to the active form. No ~12 kDa prodomain was visible upon KLK14 treatment, while furin released an intact profragment observed on SDS-PAGE. This KLK14-mediated prodomain processing may be vital in the disruption of the non-covalent association between the MMP14 prodomain and catalytic domain. As observed in the activity assay using a synthetic substrate, KLK14-mediated MMP14 activation was more efficient than furin, which may be due to the prodomain still being non-covalently attached serving as inhibitor of MMP14 [32]. In addition, the identified G^298^–I^299^ internal cleavage of MMP15 was located in the hinge region connecting the catalytic domain and hemopexin domain [33]. It is difficult to predict the impact of this cleavage on the protease activity of the resulting MMP15 and its specificity towards protein substrates. However, the appearance of a corresponding band in the zymogram analysis indicated ability to cleave gelatin and suggested that this specific auto-processing by initial KLK14-mediation may be responsible for shedding of the active catalytic domain of MMP15 from the cell surface. Interestingly, the cell surface localization of MMP15 was essential for its ability to cleave collagen in cell-based assays and it was required for exhibition of invasive cancer cell phenotypes [34], [35], while soluble MMP15 was found to display a nearly 13-fold higher activity on triple-helical substrates compared to the cell-surface located form [36]. This suggests, that release of the mature MMP15 from the cell surface may be an important regulatory step, resulting in enhanced triple helical collagen I degradation. Furthermore, lower concentrations of KLK14 induced step-wise MMMP16 profragment processing, as revealed by two truncated proforms, one form corresponding to KLK14 specificity (KKPR^100^↓CG) and the other to the specificity of MMP16 [23]. The processing of KKPR^100^↓CG, illustrates that KLK14 recognizes the site adjacent to cysteine-sulfhydryl group within the prodomain that associates with the catalytic zinc, which may disrupt the noncovalent interaction vital in releasing an active MMP16. Lastly, two distinct bands were present in the proMMP17 untreated samples both with identical N-termini, suggesting the difference between these products is in the C-terminal region. KLK14 processed both proMMP17 forms and generated two bands, both again with the same N-termini, three amino acids upstream at the motif TQAR^122^↓RRPQ. However, as described in the manufacturer documentation, the ProMMP17 activation site R^125^ was exchanged to P^125^ therefore the native activation site is not accessible for proteolysis, which in turn targets a secondary activation site.

The verification of the hydrolysis sites at the expected activation location (except for ProMMP17) indicates the proper processing of the proMMP by KLK14, however, it does not confirm the production of the active metalloproteinase from its zymogen form. To validate that KLK14-mediated processing does in fact release a mature and functional MMP proteinases, gelatin zymography was employed. It confirmed MMP14, 15 and 16 as functional proteinases since clear bands appeared at the expected molecular mass shifts of processed MMPs. Despite the correct mass shift, ProMMP17 displayed no gelatinolytic activity, either due to the abovementioned modification of the native activation site, or the natural low activity of MMP17 on gelatin as a substrate [24]. It is worth noting, however, that all commercially available proMMP14-17 preparations do not contain the membrane-interacting domain, located in the C-terminal region of the mature forms, which may impact activation kinetics in vivo. Therefore, demonstrating whether KLK14 activates MT-MMP in a cellular system provides increasing evidence of an inter-activating KLK and MT-MMP network.

Recently, a KLK4/KLK14 activity system on the cell surface of COS-7 cells was described [15]. The authors propose a cell surface-located organization of a proteinase complex, containing hepsin and TMPRSS2, two membrane-bound serine proteinases, which recruit KLK4 and KLK14 and activate proMMP3 and proMMP9. In this complex, KLK14 undergoes specific activation and accumulates in the active form, bound to the plasma membrane by a so far unknown mechanism. The role of cell surface-bound KLK14 in that system was not identified. However, based on the data presented in our study, we believe and propose that the proximity of MT-MMPs to membrane-attached, proteolytically active KLK14 may provide a yet undescribed platform for the activation of MT-MMPs.

As exemplified by the archetypical member of the subfamily, MT1-MMP (MMP14), this subclass of MMPs was reported to undergo furin-dependent activation in the compartments of the late secretory pathway however, a body of research challenges this earlier assumption [11]. An alternative activation pathway was identified in furin-deficient RPE.40 cells which were able to display active MMP14 with the same N-terminus as the native form found in furin-expressing COS-7 cells [11]. As furin expressing CHO-K1 cells processed the membrane-bound form to the same extent as RPE.40 cells, while also converting the soluble form of MMP14, the authors concluded, that furin is required only for the maturation of the soluble form. Other, yet unidentified enzymes are responsible for the processing of membrane bound MMP14 [11]. Similarly, the non-constitutive, furin-independent processing of proMMP14 was also reported to occur in fibrosarcoma HT-1080 and CCL-137 normal fibroblast cells, where the 63 kDa proform of MMP14 was detected in both, the tumor-derived cell line and in normal cells upon PMA stimulation [37], [38]. Thus, a furin-independent activation pathway needs to be further elucidated since active MMP14 was still present in furin-deficient cells [39]–[41].

MMP14 is expressed quite low in cells normally between 100,000 – 200,000 sites/cell which makes it difficult to perform biochemical analysis and investigate functional parameters [42]. To delve into its processing, the *MT1-MMP (MMP14)* gene is transfected either transiently or stably which results in MMP14 overexpression leading to a plethora of cell surface forms; 63 kDa proenzyme, 54 kDa enzyme and a 39 – 45 kDa degradation products [25]. The work presented here shows MMP14 to be processed on the cell surface by KLK14 using murine fibroblasts stably overexpressing MMP14. The ~63 and 58 kDa forms detected by immunoprecipitation are consistent with previous reports and were identified as proMMP14 and active MMP14, respectively [25], [26]. Indeed, the intensity of the 58 kDa band increased upon KLK14 treatment. In addition, a lower-molecular weight MMP14 form of around 56 kDa appeared. Intriguingly, this may present an alternative location of processing the membrane anchored proMMP14 by KLK14, as potential locations corresponding with KLK14 specificity are present ~1 and 4 kDa downstream in the MMP14 sequence (TPK^134^ and IRK^146^, respectively). Furthermore, in our experiments furin did not process cell surface proMMP14, also consistently with previous reports, indicating that furin acts exclusively in the Golgi apparatus and/or on soluble, not-membrane-bound forms of MMP14 [39].

Researchers confirmed that there is a 2-step activation process for MT1-MMP. The linkage (RRPRC^93^GVPD) in the prodomain maintains the latent proenzyme by chelating the cysteine-sulfhydryl group to the active site zinc, which furin hydrolysis alone is not enough to disrupt the interaction [32]. First, an MMP-dependent cleavage at the bait region site (PGD↓L^50^) is required and initiates the liberation of MMP14. This results in an activation intermediate in which furin or another proprotein convertase then hydrolyzes the RRKR^111^↓Y^112^ sequence providing an active form to the membrane [43]. In addition to this two-step mechanism, another MMP cleavage site (GAE?I^103^) was identified which results in a 9 residue longer N-terminus of active MMP14 with the RRKR^111^↓Y^112^ sequence still remaining. Most importantly, mutant forms of the MMP dependent cleavage site (PGD↓L^50^) and furin activation site (RRKR^111^↓Y^112^) sequences in HT 1080 cells showed both proenzyme and enzyme forms of MMP14 on their cell surface, thus again confirming a furin-independent pathway of MMP14 processing [32]. This cell-surface processing sheds more light on the potential of various secreted MMP14 forms to be differentially regulated in normal and cancer cells by extracellular KLK14. In addition to KLK14 hydrolyzing the furin recognition site (RRKR^111^↓Y^112^), KLK14 may also recognize the trypsin-like sequence, RRPR^92^CGVPD, which is specifically adjacent to the site of the cysteine-sulfhydryl group within the prodomain that associates with the catalytic zinc in the active site. In the work presented here, when recombinant proMMP14 was incubated with furin, the released prodomain was intact and visible which confirms furin hydrolysis is limited to only the furin recognition site [43]. On the other hand, KLK14 further degraded the released prodomain since it was no longer visible at 12 kDa as compared to furin. Thus, this indicates the additional KLK14 hydrolysis in the prodomain which may be vital in the release non-covalent association of the prodomain and catalytic domain once KLK14 or furin cleave at the furin recognition site. Intriguingly, KLK14 recognized this highly conserved cysteine linkage sequence, KKPR^100^↓CGVPD, within the prodomain of recombinant MMP16. This illustrates the possibility of KLK14 to disrupt the prodomain association with the catalytic zinc in other proMMPs in vivo, providing a universal activation mechanism for the entire proMMP family by KLK14. Nonetheless, recombinant proMMP2 also contains the corresponding ^97^MRKPR^101^C sequence, yet no KLK14-mediated activation was observed, indicating that accessibility to this sequence may be selective and depend on protein conformation.

Cell surface-located, membrane type MMPs have been identified as important players in bone remodeling as well as in wound healing, growth factor signaling, and in immune functions [44]–[46], while KLKs are implicated in immune functions and TGFβ activation [47]. Interestingly, the presence of both MT-MMPs and KLKs was identified in many types of cancer [46], [48]. A body of research indicates MMP14 as the main membrane type MMP essential for cancer cell escape and hence, tumor progression [49], [50], which is partially attributed to proMMP2 activation, but also for the ability of MMP14 to directly degrade type I collagen and to promote cellular invasion in 3D collagen matrices [51]. Similarly, MMP15 and MMP16 were shown to promote invasiveness of tumor cells in 3D fibrin matrices [10], [52], while in particular MMP16 is highly expressed in aggressive melanoma [10]. MMP17 was implicated as a protease critical for breast cancer metastasis in animal models [53]. KLK14 was similarly implicated in many human tumors. Most notably, KLK14 levels were found to predict poor outcomes in prostate cancer patients [54] and elevated levels of both, transcript and protein, were found in malignant breast cancer tissues [55], [56]. In correlation, increased levels of MMP14 were detected in prostate cancer cells [57], [58] and active MMP2 was found to be inversely correlated with the disease-free survival time in prostate cancer patients [57]. Furthermore, elevated levels of MMP14 were associated with poor prognosis and invasiveness in breast cancer [59], [60]. Recently, the involvement of MMP14 was also indicated in epithelial-to-mesenchymal transition in squamous cell carcinoma [61], [62] and prostate cancer alike [63].

It is therefore exciting to speculate, that the herein described potential of KLK14 to activate membrane-type MMPs may provide a novel mechanism facilitating tumorigenesis, initial cancer cell invasion and subsequent tumor progression at the sites of metastases formation.

## EXPERIMENTAL PROCEDURE

### Cloning of HmuY-based CleavEx fusion proteins

The CleavEx library was constructed based on a proteolysis-resistant *Porphyromonas gingivalis* HmuY protein (accession number ABL74281.1) as a carrier, *via* PCR cloning. Firstly, the HmuY gene was amplified using primers forward: 5’–atatgcggccgcagacgagccgaaccaaccctcca–3’ and reverse: 5’–atactcgagttatttaacggggtatgtataagcgaaagtga–3’ from whole-genomic DNA isolated from *P. gingivalis* strain W83. PCR was conducted for 35 cycles with initial denaturation at 98°C, followed by 40s annealing at 68°C and 30s extension at 72°C, using Phusion DNA polymerase (Thermo Scientific) and T100 Thermal Cycler (Bio-Rad). The HmuY PCR product was further amplified in three consecutive PCR reactions with primers specific to the 5’ HmuY fragment and a 3’ specific primer introducing additional nucleotides dependent on the designed sequence (Table 3) at the same conditions. All sequences were designed based of the accession number from the Uniprot database (www.uniprot.org); MMP1 (P03956), MMP2 (P08253), MMP3 (P08254), MMP7 (P09237), MMP8 (P22894), MMP9 (P14780), MMP10 (P09238), MMP11 (P24347), MMP12 (P39900), MMP13 (P45452), MMP14 (P50281), MMP15 (P51511), MMP16 (P51512), MMP17 (Q9ULZ9), MMP19 (Q99542), MMP20 (O60882), MMP21 (Q8N119), MMP23 (O75900), MMP24 (Q9Y5R2), MMP25 (Q9NPA2), MMP26 (Q9NRE1), MMP27 (Q9H306) and MMP28 (Q9H239). Lastly, the final PCR reaction was ligated into a modified pETDuet plasmid, according to the manufacturers protocol, with potential tryptic cleavage sites removed from the MCS using QuickChange (Agilent Technologies). An alternative method was also used for the fusion protein-encoding sequences by using Phusion Site-Directed Mutagenesis (Thermo Scientific) *via* sequence exchange from a previously prepared CleavEx construct (Table 4). The final product was transformed into competent *E. coli* T10 cells then purified and sequenced. All library DNA sequences were identified to be as intended.

**Table 3:**
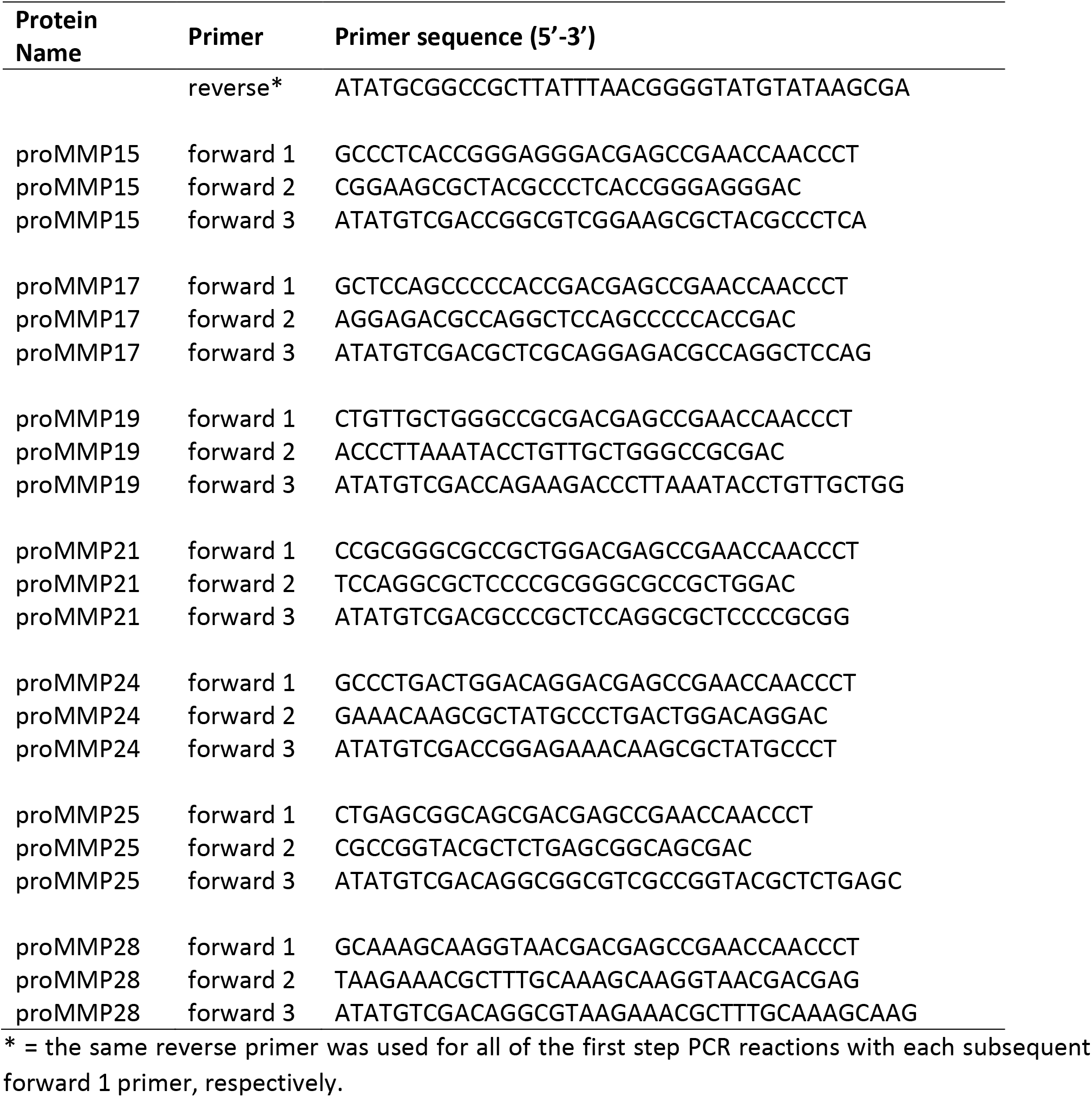
Primers used for generating the proMMP library using three consecutive PCRs.

**Table 4:**
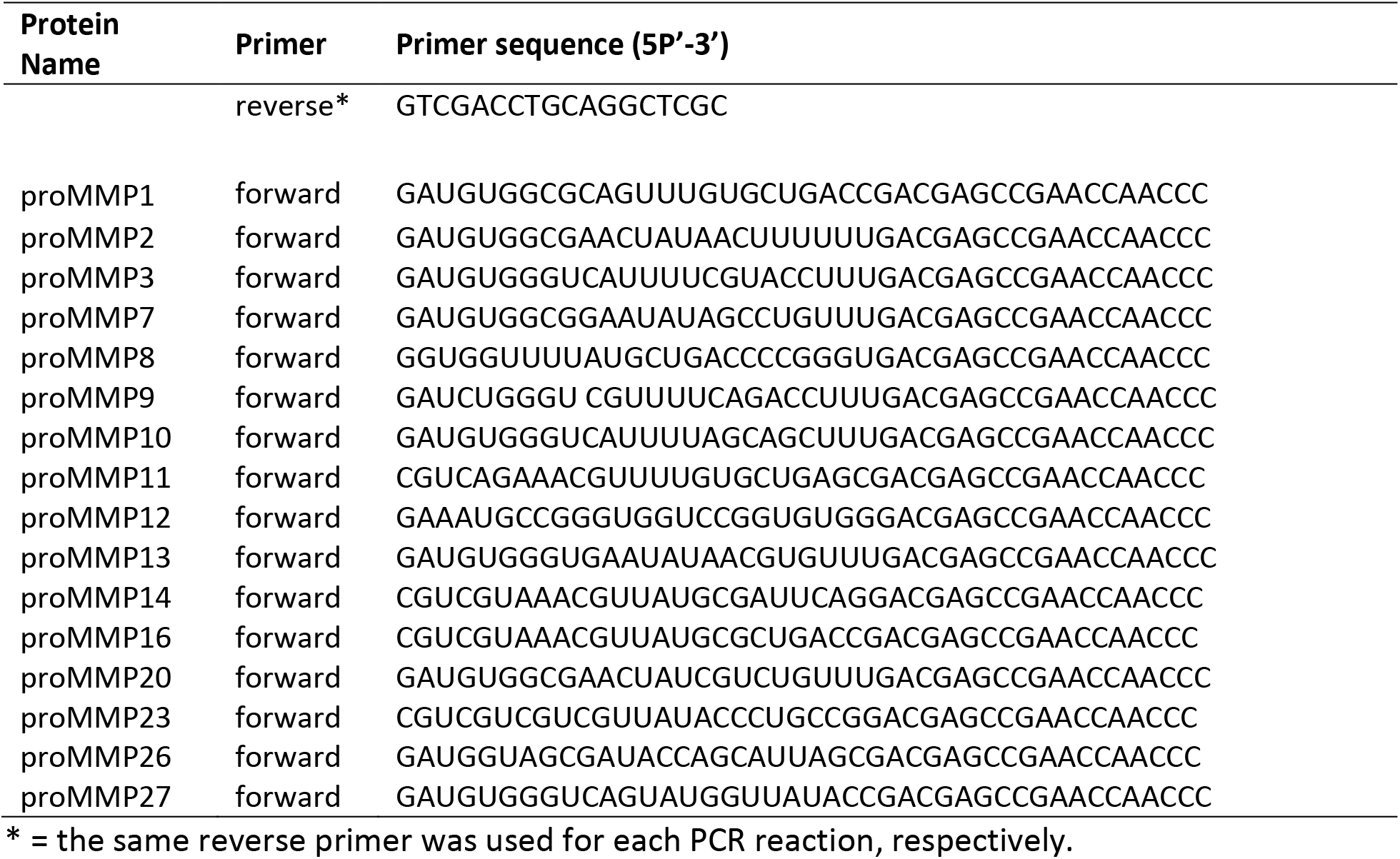
Primers used for generating the proMMP library using mutagenesis.

### Expression and purification of CleavEx library fusion proteins

The library of the designed fusion proteins covering proMMP activation sequences were expressed using an *E. coli* BL21 expression system. Following the 0.5mM IPTG induction at OD_600nm_ = 0.5-0.6, the bacterial culture protein production was facilitated for 3hrs at 37°C with shaking. Then, the bacteria were spun down and the pellet was suspended in buffer A (10 mM sodium phosphate, 500 mM NaCl and 5 mM imidazole, pH 7.4) and sonicated (15 min at 16°C, pulse 6s, amplitude 70%). Supernatant of the soluble proteins was then loaded onto the HisTrap™ Excel (GE Healthcare) column in buffer A and eluted with a linear gradient of 0-100% of 1 M imidazole in buffer A in 20 column volumes (CV). Protein containing fractions were pooled together and exchanged into 50 mM Tris pH 7.5 then purified by ion exchange chromatography using a MonoQ 4.6/100 PE column (GE Healthcare) with a linear gradient of 0-100% 50 mM Tris pH 7.5, 1 M NaCl in 15 CV. Purity of all the library products was verified by SDS-PAGE.

### Expression and production of KLK14

The gene encoding human proKLK14 was custom-synthesized by Life Technologies with a codon usage optimized for *Leishmania tarentolae* and cloned into the pLEXSY_I-blecherry3 plasmid (Cat. No. EGE-243, JenaBioscience) using NotI and XbaI restriction sites. All preparations for transfection, selection and expression in host strain T7-TR of *L. tarentolae* were performed according to the JenaBioscience protocol for inducible expression of recombinant proteins secreted to the medium. Expression of proKLK14 was induced with 15 μg/ml of tetracycline (BioShop) and was carried out for 3 days. The protein of interest was purified from the medium on HisTrap™ Excel (GE Healthcare) as described above. Obtained fractions were analyzed by SDS PAGE and fractions containing proKLK14 were concentrated with Vivaspin^®^ 2 (Sartorius) and further purified on Superdex s75 pg (GE Healthcare) in 20 mM Tris pH 7.5, 0.5 M NaCl. After purification and self-activation in 37°C for 24hrs, KLK14 was active-site titrated as described in Kantyka *et al*. [64].

### Screening the CleavEx_proMMP_ library with KLK14 and N-terminal sequencing of KLK14-released fragments

CleavEx proteins were incubated at a 1:1000 and 1:200 KLK14:CleavEx_proMMP_ molar ratio, corresponding to 50 and 250 nM KLK14, respectively in 50 mM Tris pH 7.5. Samples were incubated at 37°C for 1 hour and after which time the reactions were immediately stopped by the addition of 50 mM DTT-supplemented SDS sample buffer (1:1) and boiled. The obtained samples were resolved using 10% Tricine SDS-PAGE. The proteins were electrotransferred onto a PVDF membrane (Amersham^TM^ Hybond^TM^, GE Healthcare) in 25 mM Tris, 190 mM glycine and 20% methanol at 100 V for 1 hour in 4°C. The membranes were blocked with 5% skim milk in TTBS (50 mM Tris-HCl, 500 mM NaCl, 0.05% Tween-20, pH 7.5) and incubated with an anti-Histag-HRP antibody (catalog no. A7058, Sigma-Aldrich) diluted 1:20 000 in TTBS. The Western blots were developed with Pierce^®^ ECL Western blotting substrate (Thermo Scientific) using Medical X-Ray-Film Blue (Agfa).

Furthermore, each hydrolyzed CleavEx_proMMP_ (1 μg) was incubated with 250 nM KLK14 for 1 hour at 37°C, the reactions were stopped and the products were resolved on SDS-PAGE and electrotransferred onto a PVDF membrane [65]. The membranes were stained with Coomassie Brilliant Blue R-250 (BioShop) and the bands of interest were sequenced *via* automated Edman degradation using a PPSQ/31B protein sequencer (Shimadzu Biotech) equipped with an LC-20AT HPLC, CTO-20A column heater and SPD20A UV detector (Shimadzu Biotech) for on-line PTH analysis. Data was recorded using proprietary software (Shimadzu Biotech) and the sequence was determined by visual inspection of the UV 269 nm chromatograms.

### SDS-PAGE analysis of KLK14-mediated recombinant proMMP processing

A total of 0.5 μg native proMMP2 (catalog no. 902-MP-010, R&D systems), 0.5 μg proMMP14 (catalog no. 918-MP-010, R&D systems), 1 μg proMMP15 (catalog no. 916-MP-010, R&D systems), 0.5 μg ProMMP16 (catalog no. 1785-MP-010, R&D systems) and 1 μg ProMMP17 (catalog no. 7796-MP-010, R&D systems) were separately incubated in 10 μl with a range of KLK14 concentrations (25-250 nM, with molar ratios from around 1:65 to 1:10 KLK14:MMP) in the presence of 5 μM batimastat (Sigma) for 1 hour at 37°C in PBS. As a positive control, a total of 1 μg native proMMP2 and 0.5 μg proMMP14 were separately incubated in 10 μl with a range of furin (catalog no. 1503-SE-010, R&D systems) concentrations (0 – 250 nM, molar ratios from around 1:65 to 1:6). The reactions were stopped with the addition of 50 mM DTT-supplemented SDS sample buffer (1:1, v:v) and boiled. The samples were resolved using SDS-PAGE as described above and then visualized with Coomassie Brilliant Blue G-250 (Bioshop) staining. Additionally, 50 nM KLK14 (proMMP14) or 100 nM KLK14 (proMMP15, ProMMP16 and ProMMP17) was incubated with each respective proMMP (0.5 μg proMMP14 and 16, 1 μg proMMP15 and 17) in the presence of 5 μM batimastat for specified periods of time (0-180 min). The final molar ratio for KLK14:MMP14 was 1:32, KLK14:MMP15 was 1:16, KLK14:MMP16 was 1:16 and KLK14:MMP17 was 1:18. The reaction in each sample was stopped as above and SDS-PAGE separation was visualized using CBB G-250.

### N-terminal sequencing of KLK14 processed recombinant proMMPs

Each native proMMP (2 μg) was incubated with 50 nM (proMMP14) or 250 nM (proMMP15, ProMMP16, ProMMP17) KLK14 for 1 hour at 37°C and resolved by SDS-PAGE as described above. Proteins were then electrotransferred onto a PVDF membrane in 10 mM CAPS, 10% methanol, pH 11 using the Trans-blot semi-dry transfer cell (Bio-Rad) at 15V for 30 min. The membrane was stained with Coomassie Brilliant Blue R-250 (BioShop) and bands of interest were sequenced *via* Edman degradation as described above.

### Zymogram analysis of recombinant proMMP by KLK14

Each native proMMP (0.5 μg) was incubated with 50 and 100 nM KLK14 (proMMP14) or 100 and 250 nM KLK14 (proMMP15, 16 and 17) for 1 hour at 37°C. After incubation, KLK14 was inhibited with 10 μM biotin-Tyr-Gly-Pro-Arg-CMK, a specific KLK14 inhibitor, and 1 μM serine protease inhibitor Kazal-type 6 (SPINK6) [66] for 15 minutes at 37°C. Next, non-reducing sample buffer (1:1) was added and samples were incubated for 30 minutes at 37°C. Then samples were separated using a Tricine-SDS gel containing 0.1% gelatin at 4°C and 40 mA. Subsequently, the gels were washed three times with 2.5% Triton X-100 followed by overnight incubation in assay buffer as recommended by the manufacturer’s protocol for each proMMP with the presence of the KLK14 inhibitors at 37°C. The next day, the zymogram was fixed by using 30% methanol with 10% acetic acid for 2 minutes and stained with 0.1% amido black in 10% acetic acid for 2 hours at room temperature followed by destaining with 10% acetic acid.

### Functional activation of recombinant proMMP14 using a synthetic substrate

ProMMP14 (10 nM) was separately incubated with KLK14 (3 nM) and furin (3 nM) in the presence of the fluorogenic substrate Mca-KPLGL-Dpa-AR-NH2 (Catalog no. ES010, R&D systems). The final substrate concentration was 10 μM in 50 mM Tris, 3 mM CaCl2, 1 μM ZnCl2, pH 8.5. Hydrolysis was recorded for 180 minutes at 37°C (λ_ex_320 nm; λe_m_405nm) using a microplate fluorescence reader (Molecular Devices Spectra Max GEMINI EM). The measurement was performed in triplicates and the initial velocities were determined via the built-in linear regression algorithm. The velocities were plotted in triplicates as mean ± SEM using GraphPad using time points 0, 5, 15, 30, 60, 90, 120, 150 and 180 minutes.

### Cell surface processing of proMMP14 (MT1-MMP) by KLK14

*Timp* 2-/- murine epithelial fibroblast (MEF) cells stably expressing FLAG-tagged MT1 – MMP transfected with the pGW1GH/hMT1-MMP expression vector [67] were seeded in a 12 – well plate (250,000 cells per well) and cultured in 1.5 ml selection medium (DMEM (Gibco) with 10% FBS (Gibco), 25 μg/ml mycophenolic acid (Sigma), 250 μg/ml xanthine (Sigma) and 1X HT supplement (Life Technologies) for 2 days at 37°C, 5% CO_2_ to 80% confluency. The selection media was aspirated and cells were gently washed with PBS, then treated with 250 μl DMEM containing either 50 nM KLK14, 50 nM furin and each enzyme preincubated (15 min at 37°C) with its specific inhibitor: either 250 nM SPINK6 [66] or 1 μM decanoyl-RVKR-CMK (Bachem), respectively and further incubated for 30 min at 37°C, 5% CO_2_.

The supernatant was aspirated and the cells were rinsed with cold PBS three times and then treated with 0.1 mg/ml EZ-link^TM^ Sulfo-NHS-LC-Biotin (Thermo Scientific) for 1 hour. Next, the cells were washed three times with PBS containing 100 mM glycine and then lysed with 500 μl RIPA buffer (10 mM Tris pH 7.5, 150 mM NaCl, 1% NP-40, 0.1% SDS, 1% DOC, 2 mM EDTA) containing 25X cOmplete™ EDTA-free Protease Inhibitor Cocktail (Roche) and 5 mM EDTA. Cells were detached and spun down 16,000 rcf for 15 min at 4°C. The protein concentration of the lysate was determined using the Pierce™ BCA Protein Assay Kit (Thermo Scientific) according to the manufacturer’s instructions. A total of 500 μg was loaded onto PureProteome streptavidin magnetic beads (Millipore) and incubated overnight at 4°C. The beads were washed twice in RIPA buffer followed by a twice wash in PBS and then suspended in 50 mM tris pH 7.5 and 6x reducing sample buffer to a final 3x reducing sample buffer (v/v) in a final 40 μl volume. Samples were boiled at 95°C for 5 minutes and resolved on SDS-PAGE and electrotransferred onto a PVDF membrane [65].

The membrane was subsequently blocked with 5% skim milk in TTBS for 2 hours at 37°C. Next, primary antibody rabbit-anti-MMP14 (catalog no. MA5-32076, Thermo Scientific) in 5% milk was added at 1:1000 overnight at 4°C. The following day, the membrane was washed four times with TTBS and secondary antibody goat-anti rabbit-HRP (catalog no. A7058, Sigma) was added at 1:60,000 in 5% milk for 2 hours at RT. Lastly, the membrane was rinsed four times with TTBS and developed with the SuperSignal^®^ West Femto Maximum Sensitivity Substrate (Thermo Scientific) using Medical X-Ray-Film Blue (Agfa).

## FOOTNOTES

This project was supported by the National Science Centre grant UMO-2013/08/W/NZ1/00696 awarded to JP. KF acknowledges support by the National Science Centre grant UMO-2016/23/N/NZ1/01517. TK acknowledges support by the National Science Centre grant UMO-2016/22/E/NZ5/00332. KBr acknowledges support by the National Science Centre grant ID 211744, Ref. 2013/08/W/NZ1/00696.

The abbreviations used are: KLK, kallikrein-related peptidase; MMP, matrix metalloproteinase; ECM, extracellular matrix; CleavEx, cleavage of exposed amino acid sequences; MT-MMP, membrane-type matrix metalloproteinase; ADAM, a disintegrin and metalloproteinase; TMPRSS2, transmembrane serine protease 2; HmuY, heme-binding protein

## References

[1] C. Frantz, K. M. Stewart, and V. M. Weaver, “The extracellular matrix at a glance.,” J. Cell Sci., vol. 123, pp. 4195–4200, 2010.

[2] N. U. B. Hansen, F. Genovese, D. J. Leeming, and M. A. Karsdal, “The importance of extracellular matrix for cell function and in vivo likeness,” Exp. Mol. Pathol., vol. 98, no. 2, pp. 286–294, 2015.

[3] D. R. Edwards, M. M. Handsley, and C. J. Pennington, “The ADAM metalloproteinases,” Mol. Aspects Med., vol. 29, no. 5, pp. 258–289, 2009.

[4] G. Sotiropoulou and G. Pampalakis, “Kallikrein-related peptidases: Bridges between immune functions and extracellular matrix degradation,” Biol. Chem., vol. 391, no. 4, pp. 321–331, 2010.

[5] A. Jabłońska-Trypuć, M. Matejczyk, and S. Rosochacki, “Matrix metalloproteinases (MMPs), the main extracellular matrix (ECM) enzymes in collagen degradation, as a target for anticancer drugs.,” J. Enzyme Inhib. Med. Chem., vol. 31, no. sup1, pp. 177–183, Nov. 2016.

[6] S. Löffek, O. Schilling, and C. W. Franzke, “Series ‘matrix metalloproteinases in lung health and disease’ edited by J. Müller-Quernheim and O. Eickelberg number 1 in this series: Biological role of matrix metalloproteinases: A critical balance,” European Respiratory Journal, vol. 38, no. 1. pp. 191–208, 01-Jul-2011.

[7] M. Seiki, “The cell surface: the stage for matrix metalloproteinase regulation of migration.,” Curr. Opin. Cell Biol., vol. 14, no. 5, pp. 624–32, Oct. 2002.

[8] M. Kajita et al., “Membrane-type 1 matrix metalloproteinase cleaves CD44 and promotes cell migration,” J. Cell Biol., vol. 153, no. 5, pp. 893–904, May 2001.

[9] P. Sena et al., “Matrix metalloproteinases 15 and 19 are stromal regulators of colorectal cancer development from the early stages,” Int. J. Oncol., vol. 41, no. 1, pp. 260–266, 2012.

[10] O. Tatti et al., “MMP16 mediates a proteolytic switch to promote cell-cell adhesion, collagen alignment, and lymphatic invasion in melanoma,” Cancer Res., vol. 75, no. 10, pp. 2083–2094, May 2015.

[11] I. Yana and S. J. Weiss, “Regulation of membrane type-1 matrix metalloproteinase activity by proprotein convertases,” Mol. Biol. Cell, vol. 11, no. 7, pp. 2387–2401, 2000.

[12] S.-T. Vilen et al., “Trypsin-2 enhances carcinoma invasion by processing tight junctions and activating ProMT1-MMP.,” Cancer Invest., vol. 30, no. 8, pp. 583–92, Oct. 2012.

[13] H.-J. Ra and W. C. Parks, “Control of matrix metalloproteinase catalytic activity.,” Matrix Biol., vol. 26, no. 8, pp. 587–96, Oct. 2007.

[14] P. Carmeliet et al., “Urokinase-generated plasmin activates matrix metalloproteinases during aneurysm formation,” Nat. Genet., vol. 17, no. 4, pp. 439–444, Dec. 1997.

[15] J. C. Reid et al., “Pericellular regulation of prostate cancer expressed kallikrein-related peptidases and matrix metalloproteinases by cell surface serine proteases.,” Am. J. Cancer Res., vol. 7, no. 11, pp. 2257–2274, 2017.

[16] M. Kalinska, U. Meyer-Hoffert, T. Kantyka, and J. Potempa, “Kallikreins - The melting pot of activity and function,” Biochimie, vol. 122. pp. 270–282, 2016.

[17] J. L. V Shaw and E. P. Diamandis, “Distribution of 15 human kallikreins in tissues and biological fluids,” Clin. Chem., vol. 53, no. 8, pp. 1423–1432, 2007.

[18] A. Page-McCaw, A. J. Ewald, and Z. Werb, “Matrix metalloproteinases and the regulation of tissue remodelling,” Nature Reviews Molecular Cell Biology, vol. 8, no. 3. pp. 221–233, Mar-2007.

[19] S. Duarte, J. Baber, T. Fujii, and A. J. Coito, “Matrix metalloproteinases in liver injury, repair and fibrosis,” Matrix Biology, vol. 44–46. Elsevier, pp. 147–156, 01-May-2015.

[20] R. J. Tan and Y. Liu, “Matrix metalloproteinases in kidney homeostasis and diseases,” American Journal of Physiology - Renal Physiology, vol. 302, no. 11. pp. F1351–61, 01-Jun-2012.

[21] H. Wójtowicz et al., “Unique structure and stability of HmuY, a novel heme-binding protein of Porphyromonas gingivalis.,” PLoS Pathog., vol. 5, no. 5, p. e1000419, May 2009.

[22] T. Kang, H. Nagase, and D. Pei, “Activation of membrane-type matrix metalloproteinase 3 zymogen by the proprotein convertase furin in the trans-Golgi network.,” Cancer Res., vol. 62, no. 3, pp. 675–81, Feb. 2002.

[23] M. Kukreja et al., “High-Throughput Multiplexed Peptide-Centric Profiling Illustrates Both Substrate Cleavage Redundancy and Specificity in the MMP Family.,” Chem. Biol., vol. 22, no. 8, pp. 1122–33, Aug. 2015.

[24] H. Kolkenbrock, L. Essers, N. Ulbrich, and H. Will, “Biochemical characterization of the catalytic domain of membrane-type 4 matrix metalloproteinase.,” Biol. Chem., vol. 380, no. 9, pp. 1103–8, Sep. 1999.

[25] V. S. Golubkov, A. V Chernov, and A. Y. Strongin, “Intradomain cleavage of inhibitory prodomain is essential to protumorigenic function of membrane type-1 matrix metalloproteinase (MT1-MMP) in vivo.,” J. Biol. Chem., vol. 286, no. 39, pp. 34215–23, Sep. 2011.

[26] M. A. Chellaiah and T. Ma, “Membrane localization of membrane type 1 matrix metalloproteinase by CD44 regulates the activation of pro-matrix metalloproteinase 9 in osteoclasts.,” Biomed Res. Int., vol. 2013, p. 302392, 2013.

[27] M. Sternlicht and Z. Werb, “How Matrix metalloproteinases regulate cell behavior,” Annu. Rev. Cell Dev. Biol., vol. 17, pp. 463–516, 2009.

[28] Y. Guo, K. A. Nguyen, and J. Potempa, “Dichotomy of gingipains action as virulence factors: From cleaving substrates with the precision of a surgeon’s knife to a meat chopper-like brutal degradation of proteins,” Periodontol. 2000, vol. 54, no. 1, pp. 15–44, Oct. 2010.

[29] H. Yoon et al., “Activation profiles and regulatory cascades of the human kallikrein-related peptidases,” J. Biol. Chem., vol. 282, no. 44, pp. 31852–31864, 2007.

[30] H. Yoon et al., “Activation profiles of human kallikrein-related peptidases by proteases of the thrombostasis axis,” Protein Sci., vol. 17, no. 11, pp. 1998–2007, Nov. 2008.

[31] M. Blaber, H. Yoon, S. I. Blaber, W. Li, and I. A. Scarisbrick, “Activation profiles of human kallikrein-related peptidases by matrix metalloproteinases,” Biol. Chem., vol. 394, no. 1, pp. 137–147, 2013.

[32] V. S. Golubkov et al., “Internal cleavages of the autoinhibitory prodomain are required for membrane type 1 matrix metalloproteinase activation, although furin cleavage alone generates inactive proteinase.,” J. Biol. Chem., vol. 285, no. 36, pp. 27726–36, Sep. 2010.

[33] P. Osenkowski, S. O. Meroueh, D. Pavel, S. Mobashery, and R. Fridman, “Mutational and structural analyses of the hinge region of membrane type 1-matrix metalloproteinase and enzyme processing,” J. Biol. Chem., vol. 280, no. 28, pp. 26160–26168, Jul. 2005.

[34] K. Hotary, E. Allen, A. Punturieri, I. Yana, and S. J. Weiss, “Regulation of cell invasion and morphogenesis in a three-dimensional type I collagen matrix by membrane-type matrix metalloproteinases 1, 2, and 3,” J. Cell Biol., vol. 149, no. 6, pp. 1309–1323, Jun. 2000.

[35] I. Ota, X. Y. Li, Y. Hu, and S. J. Weiss, “Induction of a MT1-MMP and MT2-MMP-dependent basement membrane transmigration program in cancer cells by Snail 1,” Proc. Natl. Acad. Sci. U. S. A., vol. 106, no. 48, pp. 20318–20323, Dec. 2009.

[36] S. Pahwa et al., “Characterization and regulation of MT1-MMP cell surface-associated activity,” Chem. Biol. Drug Des., vol. 93, no. 6, pp. 1251–1264, Jun. 2019.

[37] P. Osenkowski, M. Toth, and R. Fridman, “Processing, shedding, and endocytosis of membrane type 1-Matrix metalloproteinase (MT1-MMP),” J. Cell. Physiol., vol. 200, no. 1, pp. 2–10, Jul. 2004.

[38] J. Lohi, K. Lehti, J. Westermarck, V. M. Kähäri, and J. Keski-Oja, “Regulation of membrane-type matrix metalloproteinase-1 expression by growth factors and phorbol 12-myristate 13-acetate,” Eur. J. Biochem., vol. 239, no. 2, pp. 239–247, 1996.

[39] J. Cao, A. Rehemtulla, W. Bahou, and S. Zucker, “Membrane type matrix metalloproteinase 1 activates pro-gelatinase A without furin cleavage of the N-terminal domain.,” J. Biol. Chem., vol. 271, no. 47, pp. 30174–80, Nov. 1996.

[40] T. Sato, T. Kondo, M. Seiki, and A. Ito, “Cell type-specific involvement of furin in membrane type 1 matrix metalloproteinase-mediated progelatinase A activation.,” Ann. N. Y. Acad. Sci., vol. 878, pp. 713–15, Jun. 1999.

[41] T. Sato, T. Kondo, T. Fujisawa, M. Seiki, and A. Ito, “Furin-independent pathway of membrane type 1-matrix metalloproteinase activation in rabbit dermal fibroblasts.,” J. Biol. Chem., vol. 274, no. 52, pp. 37280–4, Dec. 1999.

[42] A. G. Remacle, A. V Chekanov, V. S. Golubkov, A. Y. Savinov, D. V Rozanov, and A. Y. Strongin, “O-glycosylation regulates autolysis of cellular membrane type-1 matrix metalloproteinase (MT1-MMP).,” J. Biol. Chem., vol. 281, no. 25, pp. 16897–905, Jun. 2006.

[43] V. S. Golubkov et al., “Proteolysis of the membrane type-1 matrix metalloproteinase prodomain: implications for a two-step proteolytic processing and activation.,” J. Biol. Chem., vol. 282, no. 50, pp. 36283–91, Dec. 2007.

[44] C. K. Tokuhara et al., “Updating the role of matrix metalloproteinases in mineralized tissue and related diseases.,” J. Appl. Oral Sci., vol. 27, p. e20180596, Sep. 2019.

[45] M. P. Caley, V. L. C. Martins, and E. A. O’Toole, “Metalloproteinases and Wound Healing,” Adv. Wound Care, vol. 4, no. 4, pp. 225–234, Apr. 2015.

[46] S. P. Turunen, O. Tatti-Bugaeva, and K. Lehti, “Membrane-type matrix metalloproteases as diverse effectors of cancer progression.,” Biochim. Biophys. acta. Mol. cell Res., vol. 1864, no. 11 Pt A, pp. 1974–1988, Nov. 2017.

[47] G. Sotiropoulou, G. Pampalakis, and E. P. Diamandis, “Functional roles of human kallikrein-related peptidases.,” J. Biol. Chem., vol. 284, no. 48, pp. 32989–94, Nov. 2009.

[48] C. a Borgoño and E. P. Diamandis, “The emerging roles of human tissue kallikreins in cancer.,” Nat. Rev. Cancer, vol. 4, no. 11, pp. 876–90, Nov. 2004.

[49] T. Yan et al., “MMP14 regulates cell migration and invasion through epithelial-mesenchymal transition in nasopharyngeal carcinoma,” Am. J. Transl. Res., vol. 7, no. 5, pp. 950–958, Jul. 2015.

[50] P. von Nandelstadh et al., “Actin-associated protein palladin promotes tumor cell invasion by linking extracellular matrix degradation to cell cytoskeleton.,” Mol. Biol. Cell, vol. 25, no. 17, pp. 2556–70, Sep. 2014.

[51] A. M. Lowy et al., “β-catenin/Wnt signaling regulates expression of the membrane type 3 matrix metalloproteinase in gastric cancer,” Cancer Res., vol. 66, no. 9, pp. 4734–4741, May 2006.

[52] K. B. Hotary et al., “Matrix metalloproteinases (MMPs) regulate fibrin-invasive activity via MT1-MMP-dependent and -independent processes,” J. Exp. Med., vol. 195, no. 3, pp. 295–308, Feb. 2002.

[53] V. Chabottaux et al., “Membrane-type 4 matrix metalloproteinase promotes breast cancer growth and metastases,” Cancer Res., vol. 66, no. 10, pp. 5165–5172, May 2006.

[54] G. M. Yousef et al., “Differential expression of the human kallikrein gene 14 (KLK14) in normal and cancerous prostatic tissues,” Prostate, vol. 56, no. 4, pp. 287–292, Sep. 2003.

[55] F. Fritzsche et al., “Expression of human Kallikrein 14 (KLK14) in breast cancer is associated with higher tumour grades and positive nodal status,” Br. J. Cancer, vol. 94, no. 4, pp. 540–547, Feb. 2006.

[56] G. Papachristopoulou, M. Avgeris, A. Charlaftis, and A. Scorilas, “Quantitative expression analysis and study of the novel human kallikrein-related peptidase 14 gene (KLK14) in malignant and benign breast tissues,” Thromb. Haemost., vol. 105, no. 1, pp. 131–137, Jan. 2011.

[57] D. Trudel, Y. Fradet, F. Meyer, F. Harel, and B. Têtu, “Membrane-type-1 matrix metalloproteinase, matrix metalloproteinase 2, and tissue inhibitor of matrix proteinase 2 in prostate cancer: identification of patients with poor prognosis by immunohistochemistry,” Hum. Pathol., vol. 39, no. 5, pp. 731–739, May 2008.

[58] Y. Wang, Y. X. Zhang, C. Z. Kong, Z. Zhang, and Y. Y. Zhu, “Loss of P53 facilitates invasion and metastasis of prostate cancer cells,” Mol. Cell. Biochem., vol. 384, no. 1–2, pp. 121–127, Dec. 2013.

[59] Y. Li et al., “The overexpression membrane type 1 matrix metalloproteinase is associated with the progression and prognosis in breast cancer,” Am. J. Transl. Res., vol. 7, no. 1, pp. 120–127, 2015.

[60] G. Yao, P. He, L. Chen, X. Hu, F. Gu, and C. Ye, “MT1-MMP in breast cancer: induction of VEGF-C correlates with metastasis and poor prognosis.,” Cancer Cell Int., vol. 13, no. 1, p. 98, Oct. 2013.

[61] L. Pang et al., “Membrane type 1-matrix metalloproteinase induces epithelial-to-mesenchymal transition in esophageal squamous cell carcinoma: Observations from clinical and in vitro analyses,” Sci. Rep., vol. 6, Feb. 2016.

[62] C. C. Yang, L. F. Zhu, X. H. Xu, T. Y. Ning, J. H. Ye, and L. K. Liu, “Membrane Type 1 Matrix Metalloproteinase induces an epithelial to mesenchymal transition and cancer stem cell-like properties in SCC9 cells,” BMC Cancer, vol. 13, p. 171, Apr. 2013.

[63] J. Cao, C. Chiarelli, O. Richman, K. Zarrabi, P. Kozarekar, and S. Zucker, “Membrane type 1 matrix metalloproteinase induces epithelial-to-mesenchymal transition in prostate cancer,” J. Biol. Chem., vol. 283, no. 10, pp. 6232–6240, Mar. 2008.

[64] T. Kantyka et al., “Inhibition of kallikrein-related peptidases by the serine protease inhibitor of Kazal-type 6,” Peptides, vol. 32, no. 6, pp. 1187–1192, Jun. 2011.

[65] P. Matsudaira, “Sequence from picomole quantities of proteins electroblotted onto polyvinylidene difluoride membranes.,” J. Biol. Chem., vol. 262, no. 21, pp. 10035–10038, Jul. 1987.

[66] K. Plaza et al., “Gingipains of porphyromonas gingivalis affect the stability and function of serine protease inhibitor of Kazal-type 6(SPINK6), a tissue inhibitor of human kallikreins,” J. Biol. Chem., vol. 291, no. 36, pp. 18753–18764, Sep. 2016.

[67] C. J. Morrison, G. S. Butler, H. F. Bigg, C. R. Roberts, P. D. Soloway, and C. M. Overall, “Cellular Activation of MMP-2 (Gelatinase A) by MT2-MMP Occurs via a TIMP-2-independent Pathway,” J. Biol. Chem., vol. 276, no. 50, pp. 47402–47410, Dec. 2001.

